# Hyperuricemia triggers Renal Tubular Epithelial Pyroptosis by using ROS to activate the NLRP3 inflammasome

**DOI:** 10.1101/2022.03.12.484115

**Authors:** Yansheng Wu, Ruiling Li, Dongdong Li, Jiaoying Ou, Jiabao Zhou, Chuanxu Wang, Jiandong Gao

## Abstract

Renal injury resulting from hyperuricemia has gained a lot of interest. Pyroptosis refers to inflammatory cell death. The activated caspase-1 cleavage, and the pivotal protein - GSDMD could have an association with the hyperuricemic kidney lesion pathogenesis. ROS is a vital NLRP3 inflammasome antagonist in various cells. We investigated the mechanism through which ROS stimulates NLRP3 to modulate pyroptosis in renal tubular epithelial cells as well as hyperuricemic rat kidneys.

**Methods:** *In vitro* cultured renal tubular epithelial cells (NRK-52E cells) were incubated with a gradient concentration of uric acid for 24 hr to investigate the pyroptosis through flow cytometry. Next, we used the inhibitors of ROS, mitochondrial ROS, NLRP3 and Caspase-1 respectively to intervene in uric acid treated cells to analyse pyproptosis and activation of ROS- NLRP3 inflammasome signal pathway. Finally, we evaluated the mechanism of hyperuricemia triggering renal tubular epithelial pyroptosis in rat kidney tissues.

**Results:** The levels of ROS and mitochondrial ROS, the mRNA and protein expression of pyroptosis-associated factors Caspase-1 (p45, p20/10), NLRP3, and GSDMD were upregulated in uric acid, the induced NRK-52E cells as well as hyperuricemic model kidneys. The inhibition of ROS, mitochondrial ROS, NLRP3, or caspase-1 in the uric acid-induced NRK-52E cells may help in controlling pyroptosis. The expression of mRNA and protein by the cytokines IL-18 and IL-1β also increased.

**Conclusions:** Generally, hyperuricemia triggered renal tubular epithelial pyroptosis via excessive ROS to modulate NLRP3 inflammasome activation in uric acid stimulated renal tubular epithelial cells as well as the oxonic acid potassium induced hyperuricemia.

## 1 INTRODUCTION

Hyperuricemia prevalence is a global challenge. Studies suggest that hyperuricemia is closely related to multiple diseases (Braga et al., 2016; Kirca et al., 2017; Yuan et al., 2017). Hyperuricemia is likely to lead to chronic kidney disease (CKD). Uric acid concentration was an independent risk factor for early kidney failure. Besides, it was “J” type associated with CKD all-cause mortality based on the findings from a 6-year, prospective cohort study of 3885 patients suffering from CKD stages 2-4. An increase in serum uric acid (SUA) is likely to cause glomerular hypertension and arteriolosclerosis including afferent arteriolopathy and tubulointerstitial fibrosis.

Uric acid (UA) has pro-inflammatory impacts on various signaling pathways (Lu et al., 2015; Luo et al., 2016). Several studies reported that soluble uric acid (Alberts et al., 2019; Murea, 2012) and uric acid crystals resulted in sterile inflammation. This in turn established danger signaling through pattern recognition receptors, including NLRP3 inflammasome, hence triggering inflammatory caspase-1(Xiao et al., 2015). Previous studies implied that the production of reactive oxygen species (ROS) secreted by the damaged mitochondria formed the mechanism behind NLRP3 activation (Cristobal-Garcia et al., 2015). Upon the activation of caspase-1, it contributed to the secretion of inflammatory cytokines interleukin-1β (IL-1β) and interleukin-18(IL-18) as well as an inflammatory cell death. Pyroptosis is the pathway via which results in inflammation. Gasdermin D (GSDMD) is the major cause of pyroptosis (He et al., 2015; Shi et al., 2015). Activated caspase-1 and −11(or −4/5 in humans) cleave GSDMD resulting in pyroptosis. Pyroptosis is associated with the formation of ruptured pores caused by the N-terminal cleavage product of GSDMD (GSDMD-NT). GSDMD-NT binds to phosphatidylinositol phosphates and phophatidylserine restricted to the cell membrane inner leaflet and cardiolopin (Ding et al., 2016; Liu et al., 2016).

Elevated serum uric acid levels have been shown to increase renal tubular epithelial cells apoptosis, as well as impair their structure and function via oxidative stress (Yang et al., 2019). Besides, hyperuricemia-induced activation of autophagy can facilitate renal fibrogenesis by converting the renal epithelial cells into a profibrotic phenotype (Bao et al., 2018). However, further investigation regarding whether a high-level uric acid induces renal epithelial pyroptosis in hypericemic kidney lesion and hyperuricemic nephropathy is critical. Studies indicate that gout-associated uric acid crystals stimulate GSDMD, which is dispensable for cell death as well as IL-1β release (Rashidi et al., 2019). From our previous study, it was evident that the renal tubules led to inflammatory injury via NLRP3 inflammasome activation (Wu et al., 2017) besides uric acid transporter down-regulation (Wu et al., 2018), in hyperuricemic kidneys. However, the knowledge on whether uric acid result in renal tubular epithelial cell pyroptosis is scanty. In this study, we aimed to investigate the intrinsic mechanism of uric acid leading to renal tubule inflammation, to improve our understanding regarding the renal injury triggered by hyperuricemia result in novel therapy against hyperuricemia as well as uric acid nephropathy.

## 2 MATERIALS AND METHODS

### 2.1 Antibodies and reagents

Rabbit anti-rat TXNIP (ab188865), Caspase-1(ab108362), IL-18(ab191860), IL-1β (ab9722) , Caspase-1 p10 (ab179515) antibody were purchased from Abcam (Britain). Rabbit anti-rat NLRP3 (NBP2-12446SS/), ASC (NBP1-78977SS) was provided by NOVUS (United States). We obtained rabbit anti-rat caspase-1p20 (AG-20B-0042) from AdipoGen (United States). The rabbit anti-rat GSDMD (L60) (#93709) were bought from Cell Signaling Technology (CST, United States). We obtained Goat Anti-rabbit IgG/FITC (bs-0295G-FITC) and IgG-HRP (bs-0295G) from Bioss (Beijing, China). Rabbit anti-rat GAPDH (10494-1-AP) was provided by proteintech (United States). The mRNA sampling and testing kit (9767), the reverse transcription kit (RR036A), and the SYBR Premix Ex Taq TM testing kit (RR420A) were provided by the Takara Biotechnology Co. Ltd. (Japan). Uric acid (C12625) was bought from sigma (United States). The Nanjing Jiancheng Bioengineering Institute (Nanjing, China) provided the ROS testing kit (E004). MitoSOX™ Red Mitochondrial Superoxide Indicator (M36008) was purchased from Sigma (United States). BioVision (United States) sold us Z-YVAD-FMK (1140-5). MCC950 (CP-456773) was provided by Medchemexpress (United States). N-acetyl-L-cysteine (NAC, A1127) and primer synthesis service were obtained from Sangon Biotech Co., Ltd. (Shanghai, China). FLICA® 660-YVAD-FMKKF17361 was purchased from Neuromics (United States). Santa Cruz Biotechnology (United States) provided Mito-TEMPO (SC-221945). SYTOX® Green Dead Cell stain (S34860) and JC-1 kit (T3168) were provided by Invitrogen (United States).

### 2.2 Cell culture and treatment

Rats’ NRK-52E cells were bought from the Cell Resource Center, Shanghai Institute of Life Sciences, Chinese Academy of Sciences. They were maintained in RPMI-1640 medium (Invitrogen Life Technologies, 22400121), that had been supplemented with 10% fetal bovine serum (FBS; Gibco, 10099-141), 100 U/ml penicillin, as well as 100 mg/ml streptomycin (Gibco, 15144120) at 37°C in a humidified 5% CO_2_ environment. The medium was replenished daily. At 80% confluency, NRK-52E cells were incubated with a gradient concentration (0, 200, 400 and 600 μg/ml) of uric acid for 24 hr. They were later harvested and subjected to pyroptosis analysis. In cotreatment experiment, the NRK-52E cells were first exposed to 200 μg/ml uric acid media for 24 hr. The UA media was then poured and divided into three groups namely UA (addition of serum-free RPMI-1640 medium), UA+ROS inhibitor (addition of 2mM NAC) and UA + mitochondrial ROS inhibitor (addition of 50Um mito-TEMPO). In another cotreatment experiment, the uric acid-pretreated cells were also divided into three groups: UA (addition of serum-free RPMI-1640 medium), UA+NLRP3 inhibitor (addition of 7.5nM MCC950), UA + Caspase-1 inhibitor (addition of 20mM Z-YVAD-FMK). Each of the groups was incubated for 24 hr followed by subsequent analysis.

### 2.3 Cellular morphological observation and cytotoxicity assay

The NRK-52E cells were subjected to UA for 24 hr. the dynamics in their morphological characteristics were examined under an inverted phase-contrast micro-scope (ECLISE Ti2, Nikon, Japan). Cytotoxicity was established via the 4, 5-dimethylthiazolyl-2 (MTT, C009, Beyotime, China) technique based on the manufacturer’s instructions.

### 2.4 Pyroptosis analysis

The NRK-52E cells were collected and re-suspended in culture plate. The cell concentration was adjusted to 5×10^5^ cells /ml. The NRK-52E cells were incubated for 30 min with FLICA® 660-YVAD-FMK loading dye and washed thrice with ice-cold PBS. The cells were then collected and added to the flow tube. They were later lucifugal incubated for 20 min with 1 μl PI stain (SYTOX® Green Dead Cell stain) loading dye. Flow cytometry (FACSCalibur, BD Biosciences, and Rutherford, NJ, United States) set the excitation wavelength at 488nm and emission wavelength at 530nm to assess the dead cells. The flow cytometry set the excitation wavelength at 488nm and emission wavelength at 530nm to analyze the caspase-1-positive cells. The index of pyroptosis was expressed as the ratio of positively stained pytoptosic cells to the total number of the counted cells × 100% (Hu et al., 2016; Tang et al., 2019).

### 2.5 ROS detection

The NRK-52E cells and the rats’ kidney tissues suspension were incubated in PBS containing 10μM dihydroethidium for 30 min at 37°C to assess the ROS levels. They were also incubated with 5μM MitoSOXTM for 10 min at 37°C to assess the mitochondrial ROS levels. Then it was washed with PBS thrice. Flow cytometry set the excitation wavelength and emission wavelength at 485/525nm and 510/580nm respectively to analyze the total ROS and mitochondrial ROS level.

### 2.6 Mitochondrial membrane potential

Dynamics in mitochondrial membrane potential (MTP) resulting from UA were determined via a membrane-permeant JC-1 dye (5, 50, 6, 60 -tetrachloro-1, 10,3,30 -tetraethylbenzimidazol-carbocyanine iodide, Molecular Probes). JC-1 dye accumulates in mitochondria in a potential-dependent manner. In healthy cells having high DCm, JC-1 accumulates in the mitochondria as J-aggregates exhibiting red fluorescence. As the MTP declines in pyroptosis or unhealthy cells, JC-1 fails to accumulate in the mitochondria. Instead it stays in the cytosol in monomeric form exhibiting a green fluorescence (Jang et al., 2015). 24h following 200 μg/ml uric acid treatment, 5 mg/ml of JC-1 dye was added to the cells then incubated at 37°C for 30 min. The cells were then washed with PBS thrice. The alteration in NRK-52E cells was observed via flow cytometry.

### 2.7 Creation of the hyperuricemic rat model and dosage

30 male rats were divided into three groups: Control group, OA group, and Allopurinol group. Each group consisted of ten rats. Oxonic acid potassium salt (OA) was dissolved in 0.5% CMCC-Na suspension thus forming gastric juice. Allopurinol was administered 10 times. The rats were gavaged using 750mg/kg OA to produce a hyperuricemic model except for the control group for 11weeks. Besides, those belonging to the allopurinol group were gavaged using allopurinol solution for 11 weeks. All the rats were provided with ad libitum access to water.

### 2.8 Rats kidney sample collection

After drug intervention, all the rats were deprived food but not water until the following morning. They were then anaesthetized with pentobarbital sodium (30 mg/kg i.p.) before the blood samples were taken via the aorta abdominalis until all the blood is out. If the rats were not completely dead after bloodletting, an additional 200mg/kg pentobarbital sodium was injected intraperitoneally. The kidney was then quickly dissected. Partial cortex tissues from the left kidney were then frozen immediately in liquid nitrogen. The total cellular protein as well as RNA were extracted from kidney tissues. The samples were stored at −80°C for further analysis.

### 2.9 Immunofluorescence

90% ethanol was used to fix the rat kidney cortex tissues, embedded in paraffin and sectioned transversely. The samples were deparaffinized, and later subjected to antigen retrieval, spontaneous fluorescence quenching and bull bovine serum (BSA) blocking before being incubated with primary antibodies at 4°C overnight. After washing them thrice with TTBS, IgG-FITC-conjugated secondary antibody was added and stored at room temperature for 1h in the dark. The nucleus was later counterstained with DAPI and the slides visualized under the fluorescence microscope.

### 2.10 Real-time PCR

The relative levels of the target gene mRNA transcripts for controlling GAPDH were determined via quantitative real-time PCR using the specific primers. Total RNA was isolated, gDNA removed, cDNA reverse-transcription performed and SYBR green reagents premixed using commercial kits based on the manufacturer’s instructions. The primers sequences were as follows: TXNIP, forward 5’-GCTCAATCATGGTGATGTTCAAG-3’ and reverse 5’-CTTCACACACTTCCACTGTCAC-3’; NLRP3, forward 5’-CAGACCTCCAAGACCACGACTG −3’ and reverse 5’-CATCCGCAGCCA ATGAACAGAG-3’; ASC, forward 5’-TTATGGAAGAGTCTGGAGCTGTGG −3’and reverse 5’-AATGAGTGCTTGCCTGTGTTGG −3’; Caspase-1, forward 5’-AAGAAGGTGGCGCATTTCCT −3’ and reverse 5’-GACGTGTACGAGTGGGTGTT −3’; GSDMD, forward 5’-ATTGGCTCTGAATGGGATAT −3’ and reverse 5’-GAGTACGGCAAGCAGACTAAA −3’; IL-1β, forward 5’-AGGATTGCTTCCAAGCCCTTGACT −3’ and reverse 5’-ACAGCTTCTCCACAGCCACAATGA −3’; IL-18, forward 5’-CTGTCGGAGAAGGTGGTCTAC-3’and reverse 5’-TGGGAGGTGAGGGATGACT −3’; GAPDH, forward 5’-TGCACCACCAACTGCTTAG −3’ and reverse 5’-GATGCAGGGATGATGTTC −3’

All the primers’ sequences were ascertained in GenBank to prevent the inclusion of inadvertent sequence homologies. PCR amplification was carried out in triplicate: at 94°C for 5 min and 40 cycles of 94°C for 30 s, 57°C for 30 s, and 72°C for 40 s. The 2^−ΔΔCt^ method was used to determine each gene’s relative quantity of the mRNA transcripts.

### 2.11 Western blot

20 mg of frozen rat kidney tissues and NRK-52E cells were homogenized in 1 ml RIPA buffer. They were later centrifuged at 12,000 rpm/min for 10 min. The protein contents of the supernatants were determined based on the Bradford technique. The total proteins were denatured via in boiling water for 10 min. Similar quantities of total proteins were separated in a 6%–12% SDS-PAGE as well as electrophoretically transferred to a polyvinylidenedifluoride (PVDF) membrane which had been preactivated with methanol in the transferring buffer. 5% skim milk was used to block the membranes for 1 h, and then incubated overnight using specific primary antibodies at 4°C. HRP-conjugated goat anti-rabbit IgG being a secondary antibody was used to detect immunoreactive bands. The immunoreactive bands were visualized via phototope-horseradish peroxidase Western blotting detection system (Cell Signaling Technologies, Beverly, MA) as well as quantified using densitometry with Molecular Analyst (Bio-Rad Laboratories, Hercules, CA).

### 2.12 Enzyme-linked immunosorbent assay (ELISA)

The serum and rat kidney tissues’ IL-1β and IL-18 levels were determined using ELISA kits concerning the manufacturer's use protocols.

### 2.13 Statistical analysis

All the values were presented as means ±. Besides, the differences among the various groups were analysed using one-way ANOVA and subsequently multiple pair-wise comparisons were conducted using the New-man-Keuls test using SPSS 21.0 software. P values that were < 0.05 were regarded as statistically significant.

## 3 RESULTS

### 3.1 Hyperuricemia triggers renal tubular epithelial pyroptosis

To analyze how uric acid affects pyroptosis in renal tubular epithelia, we used various uric acid concentrations to stimulate rat renal tubular epithelium (NRK-52E). The renal tubular epithelial cells (RTECs) were cultured as well as treated with 0, 200, 400, 600 μg/ml uric acid for 24 hours in vitro. The cells lacking uric acid stimulation appeared round or oval. Besides, they were closely connected based on the typical morphological characteristics of paving epithelial cells. Following the stimulation of the uric acid at various concentrations for 24 hr, the cells were considerably flatter as well as irregular. Since the concentration of uric acid increased, the cells were not closely connected, the intercellular spaces enlarged as a result of pyroptosis. The rates of cell survival and inhibition were similar to the peformance under the microscope(Fig.S1). The results demonstrated that the positive pyroptosis increased significantly (*P* < 0.01 intervened with 200 μg/ml uric acid, *P* < 0.05 intervened with 400 and 600 μg/ml uric acid, respectively) in RTECs in comparison to 0 μg/ml uric acid priming(Fig.1). It was realized that 200 μg/ml uric acid-induced the most considerable tubular epithelial pyroptosis. This suggested that a higher concentration of uric acid led to pyroptosis as a result of the imbalance in intracellular as well as extracellular osmotic pressure. Besides, pyroptosis as a result of immune cells inflammation diminished. As such t, 200 μg/ml uric acid was used as the model concentration to activate renal tubular epithelial cells in the subsequent tests. The findings suggested that uric acid may trigger renal epithelial pyroptosis. However, the mechanism behind uric acid induced renal tubular epithelial pyroptosis is required to elucidate.

**Fig. 1:**
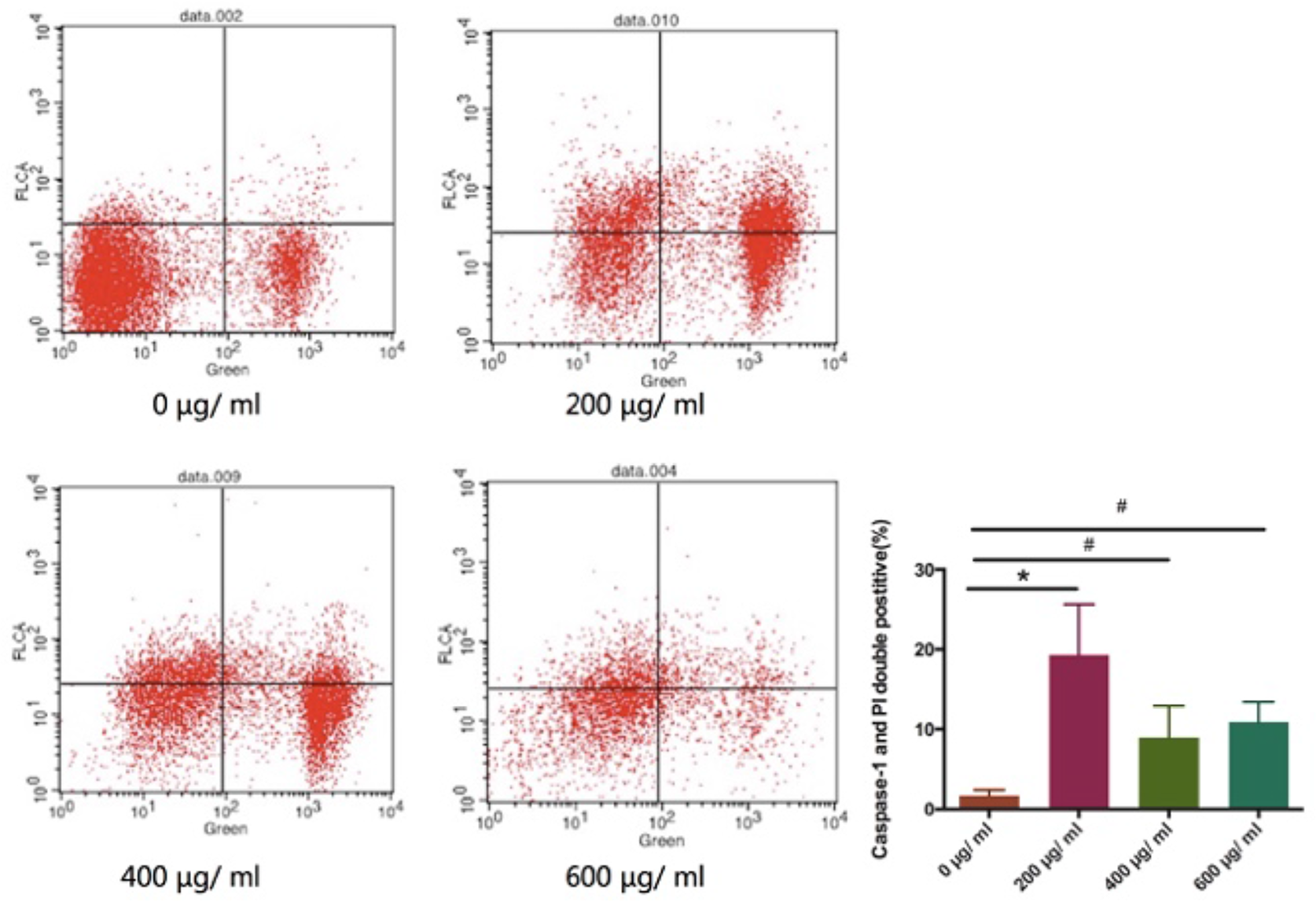
Renal tubular epithelial cells pyroptosis stimulated with various concentrations of uric acid. (A) Morphological characteristics of NRK-52E cells treated with 0, 200, 400, 600 μg/ml uric acid for 24 hours. (B)The rate of cell survival. (C) Inhibition via MTT method (n=6). (D)The pyroptosis percentage of the various concentrations of uric acid (0, 200, 400, 600 μg/ml) (n=3). The data were expressed as mean ± SD. *P < 0.01, #P < 0.05.

### 3.2 Hyperuricemia induces ROS production and inhibition of ROS reduces pyroptosis

We realized that uric acid increased the level of ROS in various kidneys’ cell lines, including the peritubular capillaries’ renal tubular cells and endothelial cells (Kang, 2018), and vascular smooth muscle cells (Tang et al., 2017), which were closely related to kidney inflammation. Also, we realized that ROS, particularly mitochondrial ROS (mtROS), triggered NLRP3 inflammasome activation in the kidneys of hyperuricemic lab rats (Wu et al., 2017). As such, we detected the production of ROS in NRK-52E to explore the association between ROS and pyroptosis. The RTECs were activated for 24 hr with 200 μg/ml uric acid. Besides, the ROS inhibitors NAC and mtROS inhibitors mt-TEMPO intervened. We detected intracellular ROS as well as mtROS levels by the fluorescence of DCFH-DA and MitoSOX probes by flow cytometry (Fig.2A-B). The ROS and mtROS production in the model group increased significantly (*P*<0.0001). In addition ROS inhibitors and mtROS inhibitors ROS and mtROS levels (*P* < 0.0001). Besides, we determined the dynamics in mitochondrial transmembrane potential (MTP) of RTECs using JC-1 kits(Fig.2C). The fluorescence ratio of the JC-1 probe multimer to monomeric was detected using flow cytometry. It was realized that the dynamics in MTP in the model group decreased significantly (*P* < 0.0001). The two ROS inhibitor groups increased the changes in MTP (*P* < 0.0001). These findings demonstrated that high levels of uric acid stimulated RTECs to induce intracellular mitochondrial injury and functional decline. As a result, ROS production increased resulting in oxidative stress as well as inflammatory damage. To ascertain the association between uric acid-stimulated ROS production and pyroptosis in RTECs, we added ROS inhibitors and mtROS inhibitors to the cultured cells respectively, before testing for pyroptosis(Fig.2D). It was also realized that the number of pyroptotic cells in the model group increased considerably (*P* < 0.0001). Both ROS and mtROS inhibition groups reduced pyroptosis (*P* < 0.0001). Besides, considering the dynamics in MPT and mtROS in uric acid-treated renal tubular epithelial cells, we can conclude that mitochondria played a vital role in cell pyroptosis.

**Fig. 2:**
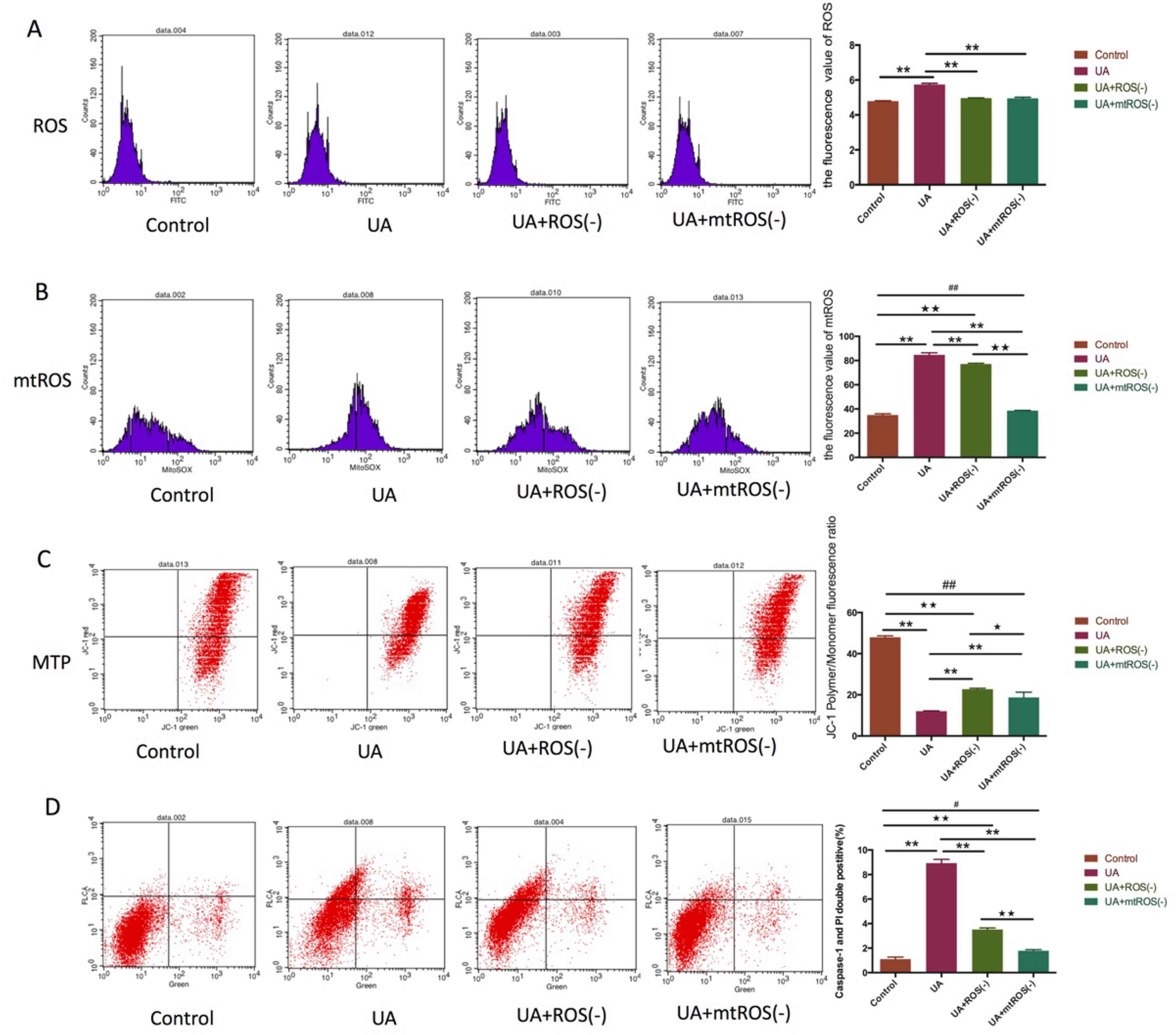
The effect of ROS inhibition on mitochondrial injury and renal tubular epithelial cell pyroptosis. (A) ROS level, (B) mitochondrial ROS level, (C) the changes in mitochondrial transmembrane potential, and (D) pyroptosis in NRK-52E cells when stimulated with 200 μg/ml uric acid or not for 24 hr via flow cytometry method (n=3). The cells were categorized into four groups: the control group (0 μg/ml uric acid), the UA group (200 μg/ml uric), the UA+ROS (−) group (200 μg/ml uric + the ROS inhibitors NAC), the UA+mtROS (−) group (200 μg/ml uric+ mtROS inhibitors mt-TEMPO). Data were expressed as mean ± SD. Compared with the UA group, \*\**P* < 0.0001, \**P* < 0.05. Compared with the UA+ROS (-) group, ⋆⋆*P* < 0.0001, ⋆*P* < 0.05. Compared with UA+mtROS (-) group, # #*P* < 0.0001, #P <0.05.

### 3.3 ROS activates NLRP3 inflammasome triggering pyroptosis in uric acid-treated renal tubular epithelial cells

Through the inhibition of ROS or mtROS respectively, we observed that expression of mRNA and protein of TXNIP, NLRP3, and Caspae-1 was up-regulated in uric acid-treated RTECs without ROS or mtROS inhibitors. However, the expression of these genes and proteins declined following intervention with ROS or mtROS inhibitors(Fig.3A). We also directly stimulated RTEC for 24 hr with 200 μg/ml uric acid, and intervened it with NLRP3 inhibitor MCC950 or Caspase-1 inhibitor Z-YVAD-FMK. Besides, pyroptosis was testified by Caspase-1 and PI double-positive using flow cytometry(Fig.3B). The findings demonstrated that the incidence of pyroptosis in the model group was significantly higher in comparison to that of the control group (*P* < 0.0001). Both NLRP3 inhibition and caspase-1 inhibition reduced pyroptosis in uric acid-stimulated RTECs in comparison to the model group (*P* < 0.0001). As such, we investigated the effect of NLRP3 or caspase-1 inhibition on the NLRP3-Caspase-1-GSDMD pathway. We realized that the expressions of NLRP3, ASC, caspase-1, and GSDMD mRNA in the control group, NLRP3 inhibition group and caspase-1 inhibition group decreased significantly (*P* < 0.0001) in comparison to the model group. Compared with the model group, NLRP3 inhibitors lessened NLRP3 protein expression (*P* < 0.0001). The expressions of caspase-1 and GSDMD protein were also down-regulated in the caspase-1 inhibition group (*P* < 0.05). Herein, we conclude that ROS-mediated NLRP3 inflammasome activation results in pyroptosis in uric acid-treated renal tubular epithelial cells. In addition, the activation of NLRP3 inflammasome resulted in massive production of inflammatory factors. Besides, GSDMD is another important component of the inflammasome, and the classical inflammatory pathway Caspase-1-GSDMD contributes to pyroptosis. On this basis, we searched the inflammatory impacts of NLRP3 or Caspase-1 inhibition on uric acid-treated RTECs(Fig.3C). The mRNA and protein expressions of IL-1β and IL-18, respectively, in uric acid-treated cells excluding NLRP3 or caspase-1 inhibitors increased significantly (*P* < 0.0001, *P* < 0.05). Both NLRP3 and Caspase-1 inhibitors decreased the mRNA and protein expression levels of IL-1β and IL-18 significantly (*P* < 0.0001). We therefore concluded that uric acid induces renal tubular epithelial pyroptosis when ROS activates the NLRP3-Caspase-1-GSDMD pathway, thus causing tubular and interstitial inflammation.

**Fig. 3:**
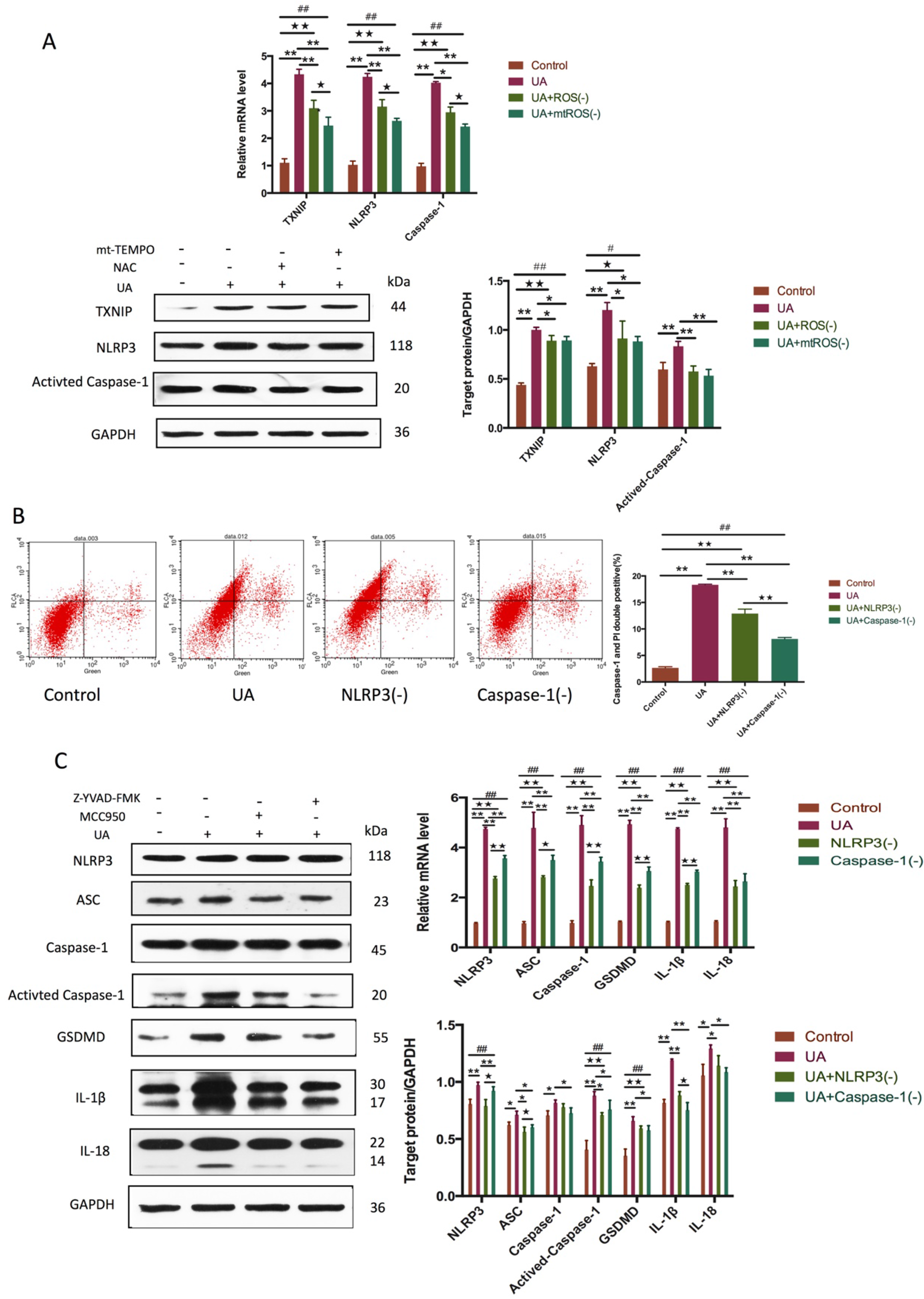
The impact of ROS and NLRP3 inflammasome on the expression of pyroptosis associated gene and protein. (A)The gene and protein expression of TXNIP, NLRP3, and Caspae-1 in NRK-52E cells under 200 μg/ml uric acid stimulation or not for 24 hr detected by Real-time PCR and western blot techniques (n=3). The cells were categorized into four groups: the control group (0 μg/ml uric acid), the UA group (200 μg/ml uric), the UA+ROS (−) group (200 μg/ml uric + the ROS inhibitors NAC), as well as the UA+mtROS (−) group (200 μg/ml uric+ mtROS inhibitors mt-TEMPO). Data were expressed as mean ± SD. Compared with the UA group, \*\**P* < 0.0001, \**P* < 0.05. Compared with the UA+ROS (−) group, ⋆⋆*P* < 0.0001, ⋆*P* < 0.05. Compared with UA+mtROS(−) group,^#^ ^#^*P* < 0.0001,^#^*P* < 0.05.(B) The pyroptosis percent in NRK-52E cells subjected to 200 μg/ml uric acid or not for 24 hr via flow cytometry method (n=3). The cells were divided into four groups: the control group (0 μg/ml uric acid), the UA group (200 μg/ml uric), the UA+NLRP3 (−) group (200 μg/ml uric + the NLRP3 inhibitors MCC950), the UA+Caspase-1(−) group (200 μg/ml uric+ Caspase-1 inhibitors Z-YVAD-FMK). Data were expressed as mean ± SD. Compared with the UA group, \*\**P* < 0.0001, \**P* < 0.05. Compared with the UA+NLRP3 (-) group, ⋆⋆*P* < 0.0001, ⋆*P* < 0.05. Compared with UA+Caspase-1(−) group,^##^*P* < 0.0001,^#^*P* < 0.05.(C) The gene and protein expression of NLRP3, ASC, Caspae-1, Activated-Caspase-1, GSDMD, IL-1β, and IL-18 in NRK-52E cells when exposed to 200 μg/ml uric acid or not for 24 hr using Real-time PCR and western blot method (n=3).

### 3.4 NLRP3 inflammasome activation in hyperuricemic rat kidneys

NLRP3 inflammasome activation including its mediated inflammatory response contribute to renal interstitial inflammation. NLRP3 inflammasome is a cytoplasmic polyprotein complex, whose final activation pathway is via reactive oxygen species (ROS) and thioredoxin interacting proteins (TXNIP) (Zhang et al., 2015). To ascertain renal tubular inflammation mechanism resulting from uric acid in vitro, we assessed the hyperuricemic rats which had been exposed to potassium oxonate. 11 weeks after modeling, serum uric acid in the model group increased significantly (*P* < 0.0001). However, both creatinine and urea nitrogen didn’t vary significantly (*P* > 0.05) (Fig.4A). Besides, consistent with the findings from in vitro experiments, ROS and mitochondrial ROS production in the model group was were increased significantly (*P* < 0.05), whereas the ROS and mitochondrial ROS levels decreased considerably in the kidneys of the treatment group (*P* < 0.05) (Fig.4B). Based on this, we speculated that oxidative stress was involved in renal injury in hyperuricemia. We also determined the role of reactive oxygen species in NLRP3 inflammasome activation, and our findings from real-time PCR and western blot analysis revealed that the genes and protein expression of TXNIP, NLRP3, ASC, Caspase-1 and Activated caspase-1 increased in the kidneys of hyperuricemic rats without allopurinol(Fig.4C). On the other hand, treatment with allopurinol reduced the expression of these genes and proteins considerably. As such, allopurinol acted as an in vivo inhibitor of the pathway we were studying. Also, we performed positioning staining of the major proteins on NLRP3 inflammasome of the renal tissues of each group via immunofluorescence. The findings demonstrated that NLRP3, ASC, and caspase-1 were extensively expressed in the renal tubular epithelial cells(Fig.4D).

**Fig. 4:**
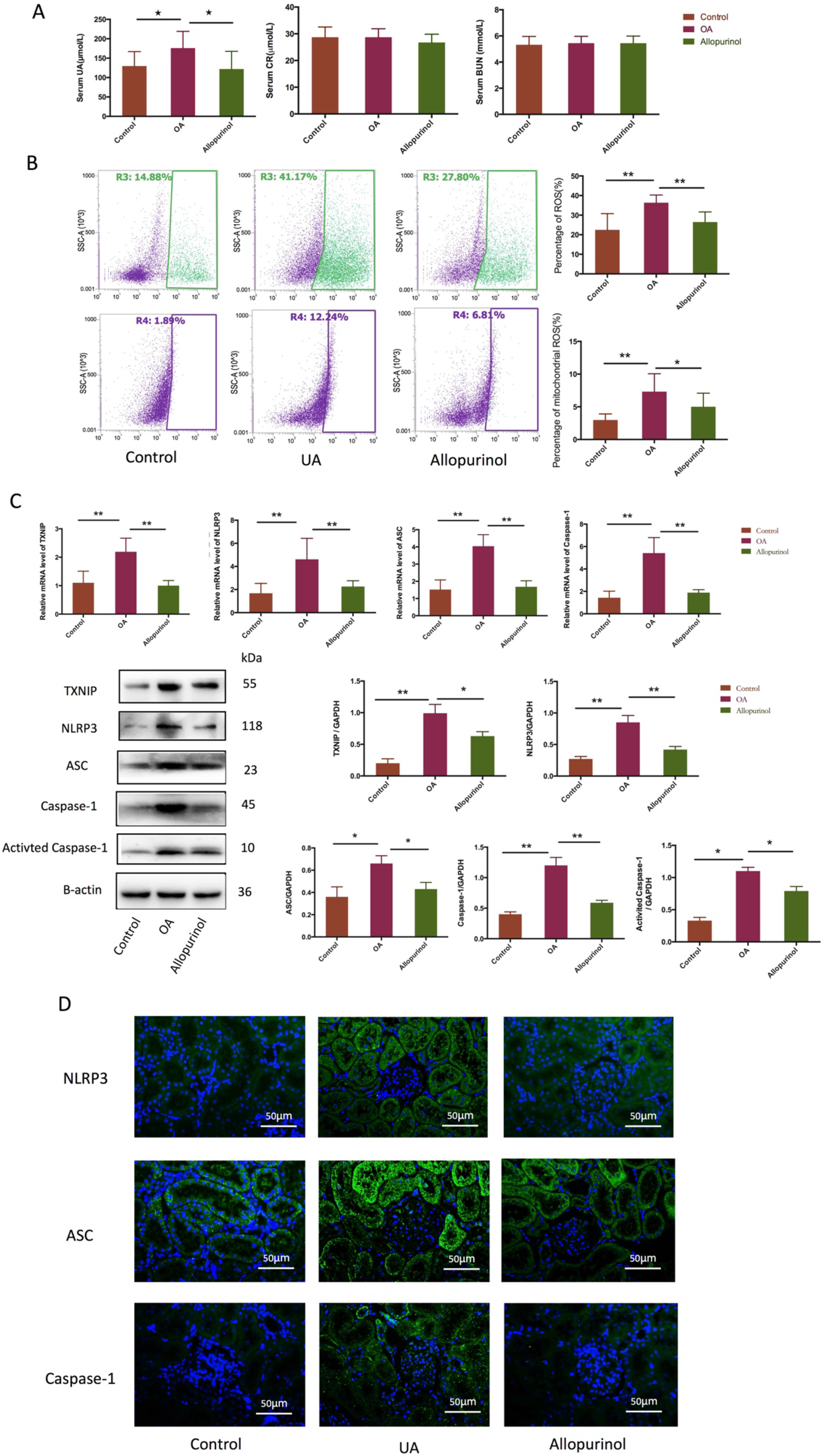
ROS activated NLRP3 inflammasome in hyperuricemic rats’ kidneys. (A) The serum uric acid, creatinine, and oxonic acid induced hyperuricemic rats (n=10). (B) ROS and mitochondrial ROS levels in renal cortical tissue cell suspension by flow cytometry method (n=10). (C)The gene and protein expression of NLRP3, ASC, Caspae-1, Activated-Caspase-1 in hyperuricemic rat kidneys (n=3). (D)Immunofluorescence of NLRP3, ASC, and Caspae-1 in hyperuricemic rat kidneys (n=3, magnification ×400). The animals were divided into three groups: the control group (n=10), the OA group (n=10) and the allopurinol group (n=10). Data were expressed as mean ± SD. \*\**P*< 0.001, \**P*< 0.05.

### 3.5 Renal tubular epithelial cells pyroptosis in hyperuricemic rats

We determined the relationship between Caspase-1 and GSDMD to understand the contribution of Caspase-1-GSDMD pathway in mediating pyroptosis as well as inflammation in hyperuricemic kidneys. We realized that the GSDMD gene and protein expression increased significantly(Fig.5A). To establish the occurrence as well as the positioning of pyroptosis in the renal tissue, we co-stained the activated Caspase-1 and GSDMD proteins via immunofluorescence(Fig.5B). Physical rupture of the cell led to the release of pro-inflammatory cytokines IL-1β and IL-18, alarmins, and endogenous danger-associated molecular patterns. As such, we tested the cytokines IL-1β and IL-18 in hyperuricemic rats’ kidneys and serum via ELISA and western blot(Fig.5C-D). We realized that the expression of IL-1β and IL-18 in the hyperuricemic model group rats’ serum and kidneys was significantly higher, suggesting a significant inflammatory response. In vivo, we confirmed the in vitro experimental findings that hyperuricemia induced pyroptosis of the renal tubular epithelial cells when ROS activated NLRP3 inflammasome, thus accelerating renal tubular inflammation.

**Fig. 5:**
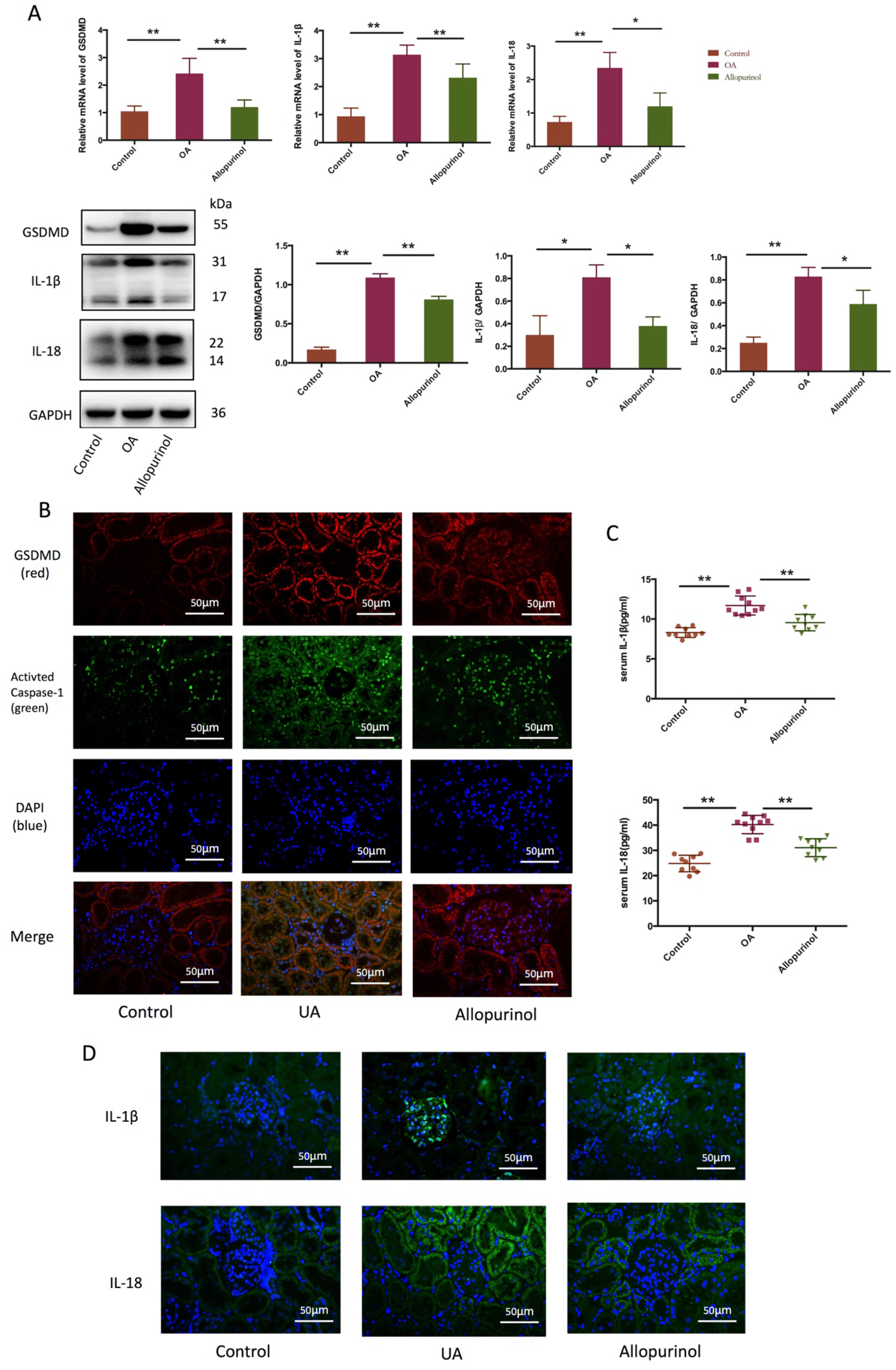
Pyroptosis and inflammatory cytokines released in the hyperuricemic rats’ kidneys. (A) The gene and protein expression of GSDMD in hyperuricemic rat kidneys (n=3). (B) Immunofluorescence co-staining of the activated Caspase-1 and GSDMD (n=3, magnification ×400). (C)The inflammatory cytokines level of IL-1β and IL-18 in hyperuricemic rats’ serum (n=10). (D)Immunofluorescence of IL-1β and IL-18 in hyperuricemic rat kidneys (n=3, magnification ×400). The animals were categorized into three groups: the control group (n=10), the OA group (n=10) and the allopurinol group (n=10). Data were expressed as mean ± SD. \*\**P* < 0.001, \**P* < 0.05.

## 4 DISCUSSION

Several studies indicated that uric acid-induced inflammatory responses were central mechanisms involving tubular injury in hyperuricemia rodents (Fan et al., 2014). The damage to the kidneys as a result of uric acid and inflammation was complementary. The impact of the high levels of uric acid on renal function was similar to that of the toxin on renal function. Hyperuricemia resulted in renal inflammation through the activation of inflammatory signaling pathways. The inflammation further contributed to renal injury via these pathways to amplify the inflammatory response (Huang et al., 2014; Neogi, 2011).

NLRP3 inflammasome activation is the critical signal pathway of renal inflammatory injury caused by high-level uric acid. ROS generation, especially from the mitochondria is regarded as the most important mechanism for the activation of the NLRP3 inflammasome/IL-1β secretion axis. Jürg Tschopp et al. suggested that mitochondria played a critical role in this process (Rock et al., 2013) and NLRP3 inflammasome was stimulated by ROS secreted from the damaged mitochondria (Zhong et al., 2016). Similarly, Shankar S. et al., observed that both ROS-dependent and non-ROS-dependent NLRP3 activation pathways contribute to mitochondrial dysfunction. Sangjun Park et al. reported mitochondrial dynamics in facilitating NLRP3 inflammasome activation, resulting in aberrant inflammation (Park et al., 2015). ROS production resulted in isolation of TXNIP from oxidized thioredoxin-1 followed by binding to the NLRP3’s LRR domain, thus activating the NLRP3 inflammasome (Zhang et al., 2015). Enhanced TXNIP led to the integration of TXNIP with NLRP3 via intracellular translocation of the TXNIP from the nucleus to mitochondria in the pathogenesis of uric acid-induced inflammatory responses (Kim et al., 2019a). In addition, the elevated serum uric acid levels can damage the renal tubular epithelial cells via oxidative stress, inflammation, increase in epithelial cells pyroptosis, as well as impair their structure and function. Mitochondria were the main organelles damaged in the process (Yang et al., 2019). However, mitochondrial damage has not been reported in uric acid-induced renal tubular epithelial pyroptosis. As such, we examined dynamics of mitochondrial membrane potential in NRK-52E cells. The findings that changes of mitochondrial membrane potential in the UA group, thus indicating severe mitochondrial injury. It is postulated that TXNIP causes mitochondria dysfunction under oxidative stress, form endogenous antagonist for thioredoxin, induce NLRP3 inflammasome activation in a ROS-NLRP3 (Kim et al., 2019b). Besides, our findings indicated that TXNIP and NLRP3 inflammasome expression was downregulated when ROS and mitochondrial ROS were inhibited in UA treated renal tubular epithelial cells. We determined ROS impacts on NLRP3 inflammasome activation as well as the impacts of NLRP3 inflammasome on pyroptosis in uric acid-treated renal tubular epithelial cells.

Both inflammation and pyroptosis are vital pathophysiological mechanisms in several diseases (Mulay et al., 2016). Scientists suggested that inflammatory necrosis is associated with an innate immune response and sterile injury can also activate the innate immune system leading to unnecessary damage in normal tissue(Rock et al., 2010). Recently (Wu et al., 2018), it was realized that the activation of NLRP3 inflammasome promoted the secretion of IL-1β and IL-18 as well as triggered a new cell death which resulted from inflammation. Pyroptosis is an example of cell inflammatory necrosis which is characterized by the formation of a ruptured pore with a diameter ranging between 1.1 and 2.4 nm in the plasma membrane. The ion concentration gradient results swelling of the cells, DNA breaks and diffuses in the nucleus, and the plasma membrane ruptures, resulting large-scale leakage of cytoplasm (Kayagaki et al., 2015). Caspase-1 and −11 (or −4/5 human) induce pyroptosis via canonical and non-canonical inflammatory pathways, respectively, to cleavage the GSDMD (Broz, 2015). Canonical inflammasome activation triggers caspase-1/GSDMD–dependent pyroptosis (Russo et al., 2016) and the inflammasome-activated gasdermin-N domain of GSDMD driving pyroptotis by selectively the inner leaflet of the plasma membrane with the pore formation activity(Qiu et al., 2017).

Caspase-1 being an effector molecule of NLRP3 inflammasome activation, the activation of ROS-NLRP3 inflammasome has proved to be considerable signal pathway on pyroptosis in various kidney diseases. Bei Zhu et al. suggested that (Zhu et al., 2020) the expression of NLRP3, cleaved-caspase1, IL-1β, GSDMD-N, the level of ROS, MDA, MCP-1, and TNF-α increased in renal tubular epithelial cells. Long noncoding RNA KCNQ1OT1 interference decreased the inflammation, oxidative stress, and pyroptosis of high glucose-induced HK-2 cells. Likewise, Chan Liu et al. reported that pyroptosis became prevalent in glucose-induced HK-2 cells due to the NLRP3 inflammation (Liu et al., 2020). The impact of high uric acid on renal tubular epithelial cell pyroptosis as well as the association between ROS and NLRP3 inflammasome in pyroptosic cells were also established

Previous studies showed that hyperuricemia resulted in kidney inflammatory lesion (Chen et al., 2019). Besides, UA triggered NRK-52E cell inflammation via NLRP3 inflammasome activation (Lu et al., 2019). In addition, hyperuricemia induced epithelial to mesenchymal transition (EMT) in the kidney. The excessive ROS production can activate multiple intracellular signaling pathways, and consequently result in excessive deposition of extracellular matrix as well as renal fibrosis. Yang-Shuo Tang et al. (Tang et al., 2019)reported that anti-oxidation could inhibit endothelial cell pyroptosis by blocking ROS/NLRP3/Caspase-1 signaling pathway. This exhibited the therapeutic effect in CKD. Besides, Orestes Foresto-Neto et al. (Foresto-Neto et al., 2018) reported that NLRP3 inflammasome inhibition ameliorates tubulointerstitial resulting from inflammation or fibrosis. Guihong Wang et al. cultured HK-2 cells that had been subjected to 100 μg/ml UA for 24 hr revealed that UA exposure inhibited cell viability, increased IL-1β and IL-18 generation in a concentration dependent manner, and activated the cleavage of gasdermin D (Wang et al., 2020). However, the mechanism on how UA induced renal tubular epithelial pyroptosis remains unclear. Our study established that the high level soluble uric acid led to renal tubular epithelial pyroptosis in vivo as well as in vitro. We found that high-level UA led to mitochondria damage as a result of excessive ROS release and NLRP3/ASC/Caspase-1-GSDMD activation. Moreover, we established that the expression of ROS, mitochondrial ROS, NLRP3 Inflammasome, GSDMD, IL-1β and IL-18 increased considerably in hyperuricemic rat kidneys.

## 5 CONCLUSIONS

Our findings established that uric acid stimulated ROS secretion in the renal tubular epithelium. Besides, the production of ROS led to the activation of the NLRP3-caspase-1-GSDMD pathway, resulting in renal tubular epithelial pyroptosis as well as tubular inflammatory lesions. Our findings further revealed that the renal tubular epithelial pyroptosis program was initiated by UA inducing NLRP3 Inflammasome activation in a mitochondrial ROS-dependent manner. It has the potential to provide molecular targets for limiting kidney injury from hyperuricemia by blocking cell pyroptosis, inflammation, or both.

## ANIMAL ETHICS

This study was conducted in accordance with safe animal testing specifications Animal experiment of this research article was approved by the experimental animal center of Shanghai University of Traditional Chinese Medicine (Safety Certificate Number: SYXK-HU-2008-0016; Animals Ethics NO.SZY201610008). The rats were raised in the experimental animal center of Shanghai University of Traditional Chinese Medicine in a 12/12 h light/dark cycle, with a feeding temperature of 26°C and relative humidity of 50%, and were given standard chow and water ad libitum during the experiment. After adaptive feeding for one week, the experimentwas started.At the end of the experiment ,all the rats were under sedation with sodium pentobarbital anesthesia through intraperitoneal injection 2%sodium pentobarbital 0.25ml/100g rat weight.

## ACKNOWLEDGEMENTS

This research was supported by the Natural Science Foundation of China (No. 81874437, No. 81904126, and No. 81373613). The authors thank Dr. Shengzhou Shan for valuable advice and support and Freescience Information Technology Co., Ltd for Language polishing.

## DATA AVAILABILITY STATEMENT

The authors promise that all the data has not been published. The related data and materials are available for sharing upon request to Jiandong Gao and Yansheng Wu.

---

**Fig. S1:**
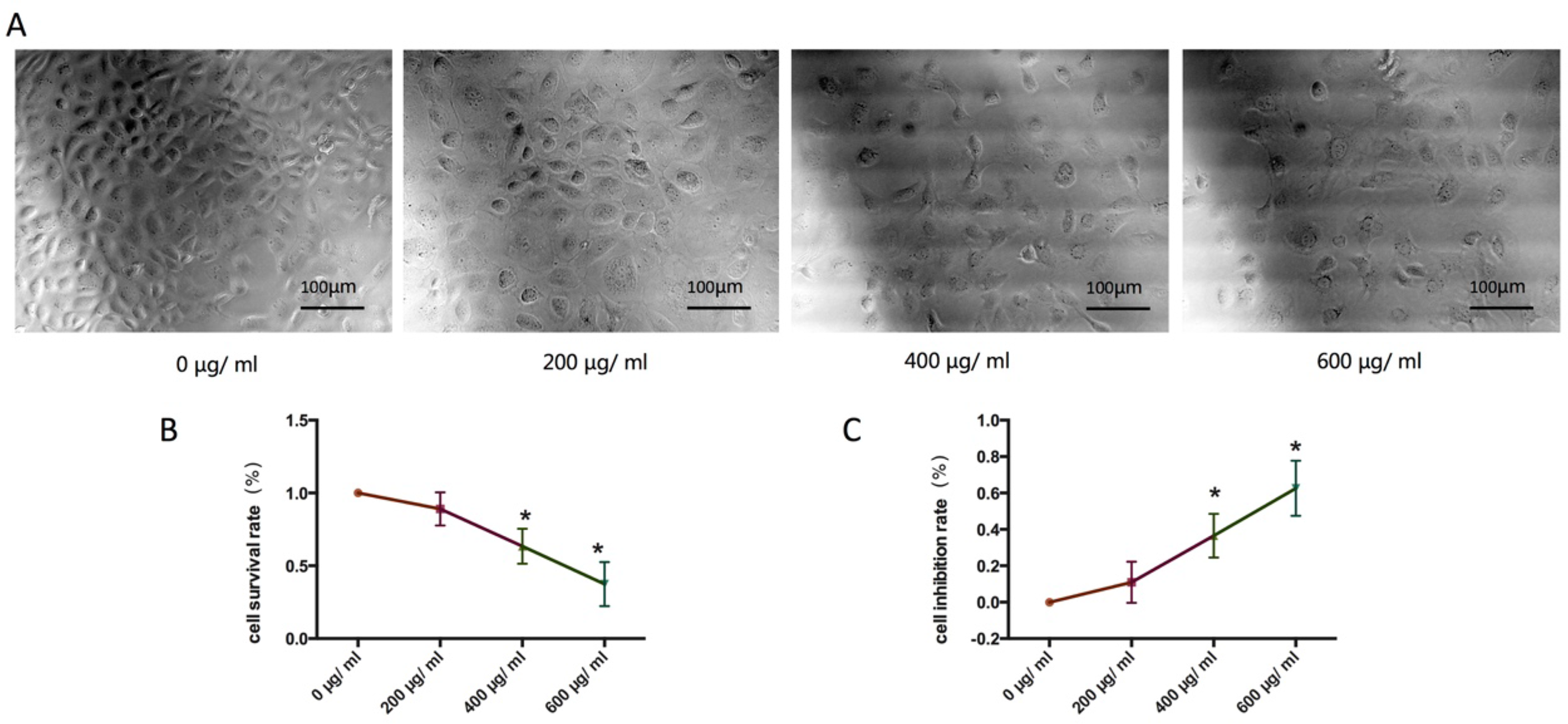
Renal tubular epithelial cells pyroptosis stimulated with various concentrations of uric acid. (A) Morphological characteristics of NRK-52E cells treated with 0, 200, 400, 600 μg/ml uric acid for 24 hours. (B)The rate of cell survival. (C) Inhibition via MTT method (n=6).

## REFERENCES

Alberts, B. M., Barber, J. S., Sacre, S. M., Davies, K. A., Ghezzi, P. and Mullen, L. M. (2019). Precipitation of Soluble Uric Acid Is Necessary for In Vitro Activation of the NLRP3 Inflammasome in Primary Human Monocytes. The Journal of Rheumatology, jrheum.180855.

Bao, J., Shi, Y., Tao, M., Liu, N., Zhuang, S. and Yuan, W. (2018). Pharmacological inhibition of autophagy by 3-MA attenuates hyperuricemic nephropathy. Clinical Science 132, 2299–2322.

Braga, F., Pasqualetti, S., Ferraro, S. and Panteghini, M. (2016). Hyperuricemia as risk factor for coronary heart disease incidence and mortality in the general population: a systematic review and meta-analysis. Clin Chem Lab Med 54, 7–15.

Broz, P. (2015). Immunology: Caspase target drives pyroptosis. Nature 526, 642–3.

Chen, Y., Li, C., Duan, S., Yuan, X., Liang, J. and Hou, S. (2019). Curcumin attenuates potassium oxonate-induced hyperuricemia and kidney inflammation in mice. Biomed Pharmacother 118, 109195.

Cristobal-Garcia, M., Garcia-Arroyo, F. E., Tapia, E., Osorio, H., Arellano-Buendia, A. S., Madero, M., Rodriguez-Iturbe, B., Pedraza-Chaverri, J., Correa, F., Zazueta, C. et al. (2015). Renal oxidative stress induced by long-term hyperuricemia alters mitochondrial function and maintains systemic hypertension. Oxid Med Cell Longev 2015, 535686.

Ding, J., Wang, K., Liu, W., She, Y., Sun, Q., Shi, J., Sun, H., Wang, D. C. and Shao, F. (2016). Pore-forming activity and structural autoinhibition of the gasdermin family. Nature 535, 111–6.

Fan, C. Y., Wang, M. X., Ge, C. X., Wang, X., Li, J. M. and Kong, L. D. (2014). Betaine supplementation protects against high-fructose-induced renal injury in rats. J Nutr Biochem 25, 353–62.

Foresto-Neto, O., Avila, V. F., Arias, S. C. A., Zambom, F. F. F., Rempel, L. C. T., Faustino, V. D., Machado, F. G., Malheiros, D., Abensur, H., Camara, N. O. S. et al. (2018). NLRP3 inflammasome inhibition ameliorates tubulointerstitial injury in the remnant kidney model. Lab Invest 98, 773–782.

He, W. T., Wan, H., Hu, L., Chen, P., Wang, X., Huang, Z., Yang, Z. H., Zhong, C. Q. and Han, J. (2015). Gasdermin D is an executor of pyroptosis and required for interleukin-1beta secretion. Cell Res 25, 1285–98.

Hu, Z., Murakami, T., Suzuki, K., Tamura, H., Reich, J., Kuwahara-Arai, K., Iba, T. and Nagaoka, I. (2016). Antimicrobial cathelicidin peptide LL-37 inhibits the pyroptosis of macrophages and improves the survival of polybacterial septic mice. Int Immunol 28, 245–53.

Huang, C. C., Lou, B. S., Hsu, F. L. and Hou, C. C. (2014). Use of urinary metabolomics to evaluate the effect of hyperuricemia on the kidney. Food Chem Toxicol 74, 35–44.

Jang, Y., Lee, A. Y., Jeong, S. H., Park, K. H., Paik, M. K., Cho, N. J., Kim, J. E. and Cho, M. H. (2015). Chlorpyrifos induces NLRP3 inflammasome and pyroptosis/apoptosis via mitochondrial oxidative stress in human keratinocyte HaCaT cells. Toxicology 338, 37–46.

Kang, D. H. (2018). Hyperuricemia and Progression of Chronic Kidney Disease: Role of Phenotype Transition of Renal Tubular and Endothelial Cells. Contrib Nephrol 192, 48–55.

Kayagaki, N., Stowe, I. B., Lee, B. L., O’Rourke, K., Anderson, K., Warming, S., Cuellar, T., Haley, B., Roose-Girma, M., Phung, Q. T. et al. (2015). Caspase-11 cleaves gasdermin D for non-canonical inflammasome signalling. Nature 526, 666–71.

Kim, S. K., Choe, J. Y. and Park, K. Y. (2019a). TXNIP-mediated nuclear factor-kappaB signaling pathway and intracellular shifting of TXNIP in uric acid-induced NLRP3 inflammasome. Biochem Biophys Res Commun 511, 725–731.

Kim, Y. G., Kim, S. M., Kim, K. P., Lee, S. H. and Moon, J. Y. (2019b). The Role of Inflammasome-Dependent and Inflammasome-Independent NLRP3 in the Kidney. Cells 8.

Kirca, M., Oguz, N., Cetin, A., Uzuner, F. and Yesilkaya, A. (2017). Uric acid stimulates proliferative pathways in vascular smooth muscle cells through the activation of p38 MAPK, p44/42 MAPK and PDGFRbeta. J Recept Signal Transduct Res 37, 167–173.

Liu, C., Zhuo, H., Ye, M. Y., Huang, G. X., Fan, M. and Huang, X. Z. (2020). LncRNA MALAT1 promoted high glucose-induced pyroptosis of renal tubular epithelial cell by sponging miR-30c targeting for NLRP3. Kaohsiung J Med Sci.

Liu, X., Zhang, Z., Ruan, J., Pan, Y., Magupalli, V. G., Wu, H. and Lieberman, J. (2016). Inflammasome-activated gasdermin D causes pyroptosis by forming membrane pores. Nature 535, 153–8.

Lu, H., Yao, H., Zou, R., Chen, X. and Xu, H. (2019). Galangin Suppresses Renal Inflammation via the Inhibition of NF-kappaB, PI3K/AKT and NLRP3 in Uric Acid Treated NRK-52E Tubular Epithelial Cells. Biomed Res Int 2019, 3018357.

Lu, W., Xu, Y., Shao, X., Gao, F., Li, Y., Hu, J., Zuo, Z., Shao, X., Zhou, L., Zhao, Y. et al. (2015). Uric Acid Produces an Inflammatory Response through Activation of NF-kappaB in the Hypothalamus: Implications for the Pathogenesis of Metabolic Disorders. Sci Rep 5, 12144.

Luo, C., Lian, X., Hong, L., Zou, J., Li, Z., Zhu, Y., Huang, T., Zhang, Y., Hu, Y., Yuan, H. et al. (2016). High Uric Acid Activates the ROS-AMPK Pathway, Impairs CD68 Expression and Inhibits OxLDL-Induced Foam-Cell Formation in a Human Monocytic Cell Line, THP-1. Cell Physiol Biochem 40, 538–548.

Mulay, S. R., Linkermann, A. and Anders, H. J. (2016). Necroinflammation in Kidney Disease. J Am Soc Nephrol 27, 27–39.

Murea, M. (2012). Advanced kidney failure and hyperuricemia. Adv Chronic Kidney Dis 19, 419–24.

Neogi, T. (2011). Clinical practice. Gout. N Engl J Med 364, 443–52.

Park, S., Won, J. H., Hwang, I., Hong, S., Lee, H. K. and Yu, J. W. (2015). Defective mitochondrial fission augments NLRP3 inflammasome activation. Sci Rep 5, 15489.

Qiu, S., Liu, J. and Xing, F. (2017). ‘Hints’ in the killer protein gasdermin D: unveiling the secrets of gasdermins driving cell death. Cell Death Differ 24, 588–596.

Rashidi, M., Simpson, D. S., Hempel, A., Frank, D., Petrie, E., Vince, A., Feltham, R., Murphy, J., Chatfield, S. M., Salvesen, G. S. et al. (2019). The Pyroptotic Cell Death Effector Gasdermin D Is Activated by Gout-Associated Uric Acid Crystals but Is Dispensable for Cell Death and IL-1β Release. The Journal of Immunology 203, 736–748.

Rock, K. L., Kataoka, H. and Lai, J. J. (2013). Uric acid as a danger signal in gout and its comorbidities. Nat Rev Rheumatol 9, 13–23.

Rock, K. L., Latz, E., Ontiveros, F. and Kono, H. (2010). The sterile inflammatory response. Annu Rev Immunol 28, 321–42.

Russo, H. M., Rathkey, J., Boyd-Tressler, A., Katsnelson, M. A., Abbott, D. W. and Dubyak, G. R. (2016). Active Caspase-1 Induces Plasma Membrane Pores That Precede Pyroptotic Lysis and Are Blocked by Lanthanides. J Immunol 197, 1353–67.

Shi, J., Zhao, Y., Wang, K., Shi, X., Wang, Y., Huang, H., Zhuang, Y., Cai, T., Wang, F. and Shao, F. (2015). Cleavage of GSDMD by inflammatory caspases determines pyroptotic cell death. Nature 526, 660–5.

Tang, L., Xu, Y. e., Wei, Y. and He, X. (2017). Uric acid induces the expression of TNF-α via the ROS-MAPK-NF-κB signaling pathway in rat vascular smooth muscle cells. Molecular Medicine Reports 16, 6928–6933.

Tang, Y. S., Zhao, Y. H., Zhong, Y., Li, X. Z., Pu, J. X., Luo, Y. C. and Zhou, Q. L. (2019). Neferine inhibits LPS-ATP-induced endothelial cell pyroptosis via regulation of ROS/NLRP3/Caspase-1 signaling pathway. Inflamm Res 68, 727–738.

Wang, G., Zuo, T. and Li, R. (2020). The mechanism of Arhalofenate in alleviating hyperuricemia―Activating PPARγ thereby reducing caspase‐1 activity. Drug Development Research 81, 859–866.

Wu, Y., He, F., Li, Y., Wang, H., Shi, L., Wan, Q., Ou, J., Zhang, X., Huang, D., Xu, L. et al. (2017). Effects of Shizhifang on NLRP3 Inflammasome Activation and Renal Tubular Injury in Hyperuricemic Rats. Evid Based Complement Alternat Med 2017, 7674240.

Wu, Y., Wang, Y., Ou, J., Wan, Q., Shi, L., Li, Y., He, F., Wang, H., He, L. and Gao, J. (2018). Effect and Mechanism of ShiZhiFang on Uric Acid Metabolism in Hyperuricemic Rats. Evid Based Complement Alternat Med 2018, 6821387.

Xiao, J., Zhang, X. L., Fu, C., Han, R., Chen, W., Lu, Y. and Ye, Z. (2015). Soluble uric acid increases NALP3 inflammasome and interleukin-1beta expression in human primary renal proximal tubule epithelial cells through the Toll-like receptor 4-mediated pathway. Int J Mol Med 35, 1347–54.

Yang, L., Chang, B., Guo, Y., Wu, X. and Liu, L. (2019). The role of oxidative stress-mediated apoptosis in the pathogenesis of uric acid nephropathy. Ren Fail 41, 616–622.

Yuan, H., Hu, Y., Zhu, Y., Zhang, Y., Luo, C., Li, Z., Wen, T., Zhuang, W., Zou, J., Hong, L. et al. (2017). Metformin ameliorates high uric acid-induced insulin resistance in skeletal muscle cells. Mol Cell Endocrinol 443, 138–145.

Zhang, X., Zhang, J. H., Chen, X. Y., Hu, Q. H., Wang, M. X., Jin, R., Zhang, Q. Y., Wang, W., Wang, R., Kang, L. L. et al. (2015). Reactive oxygen species-induced TXNIP drives fructose-mediated hepatic inflammation and lipid accumulation through NLRP3 inflammasome activation. Antioxid Redox Signal 22, 848–70.

Zhong, Z., Umemura, A., Sanchez-Lopez, E., Liang, S., Shalapour, S., Wong, J., He, F., Boassa, D., Perkins, G., Ali, S. R. et al. (2016). NF-kappaB Restricts Inflammasome Activation via Elimination of Damaged Mitochondria. Cell 164, 896–910.

Zhu, B., Cheng, X., Jiang, Y., Cheng, M., Chen, L., Bao, J. and Tang, X. (2020). Silencing of KCNQ1OT1 Decreases Oxidative Stress and Pyroptosis of Renal Tubular Epithelial Cells. Diabetes Metab Syndr Obes 13, 365–375.

